# Arg80 Orchestrates a Metabolic–Translational Proteostasis Network in S. cerevisiae

**DOI:** 10.1101/2025.08.18.670877

**Authors:** Akanksha Sharma, Shikha Rao, Zainab Zaidi, Kedar Padia, Vikramaditya Singh, Kausik Chakraborty

## Abstract

Proteostasis or protein homeostasis is essential for cellular function and organismal health. While many models of cytosolic proteostasis emphasize heat shock response as critical for regulating cellular protein folding, a comprehensive understanding of transcriptional modules that may regulate cytosolic protein folding is lacking. Through a focused screening for transcription factors, we provide evidence that a number of transcriptional programs unlinked to canonical proteotoxic response are involved in maintaining cellular homeostasis of protein folding. Among these, Arg80, a regulator of arginine metabolism, was activated during proteotoxic stress and found essential for mitigating it. We show that proteostasis imbalance in arg80Δ cells is caused by excess arginine accumulation, which is sufficient to impair protein folding. We reveal a complex interplay between arginine repression (Arg80), trehalose biosynthesis (Tps2), and the integrated stress response (Gcn2) in combating proteotoxic insults. We posit that multiple metabolic pathways integrate with classical protein quality control networks to regulate the protein folding environment. Harnessing these metabolic circuits may offer new avenues to modulate proteostasis when needed.

## INTRODUCTION

The ability of cells to maintain protein homeostasis (proteostasis) is fundamental for cellular function and organismal health. Dysregulation of proteostasis underlies diverse pathologies, including neurodegenerative diseases and aging, across eukaryotic systems (Labbadia & Morimoto, 2015; Hipp *et al*, 2019). Cellular proteostasis is frequently studied using non-physiological stressors such as elevated temperatures (Martin Vabulas *et al*, 2010; Morimoto & Santoro, 1998; Gasch *et al*, 2000) or chemical agents like Dithiothreitol (DTT) (Braakman *et al*, 1992), tunicamycin (Wu *et al*, 2018), or thapsigargin (Lindner *et al*, 2020); (Xu & Wang, 2024), which dramatically disrupt protein folding environments in specific cellular compartments, including the secretory pathway or mitochondria (Oslowski & Urano, 2011). These agents induce global misfolding, causing simultaneous perturbations to hundreds or thousands of proteins. Additionally, genetic screens designed to uncover proteostasis regulators have typically employed aggregation-prone, disease-associated proteins such as polyglutamine-expanded Huntingtin (Silva *et al*, 2011; Peters *et al*, 2018), α-synuclein (Rousseaux *et al*, 2018; Hallacli *et al*, 2022), and amyloid-β (Nair *et al*, 2014; Luu & Macreadie, 2018). While these models have been instrumental in identifying key components of the proteostasis network, they may not accurately reflect the cellular responses to more subtle or chronic proteotoxic challenges. Specifically, such stressors can activate broad stress responses due to their inherent toxicity, which might overshadow the nuanced regulatory mechanisms engaged during exclusive protein misfolding events. Therefore, there is a pressing need to explore proteostasis regulation under conditions that more closely mimic physiological stress to uncover additional pathways and factors involved in maintaining protein homeostasis.

Transcriptional responses to stress play a critical role in sensing and transmitting signals to the nucleus, ultimately activating protective cellular pathways. In yeast, transcription factors such as Hsf1 and Msn2/4 respond to stress by upregulating heat shock proteins (Hsps) and molecular chaperones to aid protein folding and prevent aggregation (Gasch *et al*, 2000; Hahn *et al*, 2004). Concurrently, genes associated with ribosome biogenesis are downregulated (Albert *et al*, 2019; Gasch *et al*, 2000), reflecting a coordinated cellular strategy to manage protein synthesis under stress. Interestingly, the transcriptional response to misfolded proteins accumulating specifically in the cytosol differs significantly from the classical heat shock response (Miyazaki *et al*, 2015). Despite these observations, the mechanistic details, such as the sensors and signaling mechanisms that mediate cytosolic unfolded protein responses (UPR), remain poorly defined. We employed a focused genetic screen in *Saccharomyces cerevisiae* to investigate these pathways using a genome-integrated, non-native misfolded protein sensor. This strategy allowed us to precisely examine cellular responses induced solely by protein misfolding, free from confounding protein functional roles.

Recent evidence highlights the critical role that metabolic intermediates play in proteostasis, functioning either as chemical chaperones (Bandyopadhyay *et al*, 2012; Dandage *et al*, 2015; Diamant *et al*, 2001) or as modulators of aggregation dynamics (Honda *et al*, 2010). Trehalose, for example, stabilizes proteins against aggregation during thermal and oxidative stresses (Tapia & Koshland, 2014; Honda *et al*, 2010; Singer & Lindquist, 1998). Arginine metabolism also influences proteostasis, displaying complex roles: it can stabilize misfolded proteins, thereby enhancing stress resistance (Ma *et al*, 2016; Baynes *et al*, 2005; Shmueli *et al*, 2019), yet can also accelerate aggregation under certain conditions (Dandage *et al*, 2015; Smirnova *et al*, 2013). Despite these important insights, the transcriptional regulators that connect metabolic pathways directly to proteostasis remain poorly characterized. To address this, we conducted a transcription factor screen for proteostasis regulators. We identified Arg80, a well-characterized transcription factor for arginine metabolism, as a key regulator of misfolded protein stress responses. Loss of Arg80 resulted in proteostasis defects, which is also mimicked in the Car1 deletion strain that is defective in arginine catabolism (Cheng *et al*, 2016); this suggests that arginine accumulation compromises cellular proteostasis, suggesting that arginine metabolism is closely linked to cellular proteostasis. Unexpectedly, we also observed upregulation of trehalose biosynthesis genes, a response typically associated with cellular stress adaptation. Further, we uncovered a functional interaction between Arg80 and Gcn2, a key regulator of translational control, linking metabolic perturbations to translational stress responses. Together, these findings expand our understanding of proteostasis regulation, demonstrating how metabolic and translational pathways intersect to maintain protein homeostasis in response to misfolding stress.

## RESULTS

### Screening of Transcription Factors that alter cellular protein folding environment

In previous studies, we utilized destabilized versions of the *Nourseothricin* (ClonNAT) resistance gene, an antibiotic resistance marker exogenous to *S. cerevisiae*, as sensors of the intracellular protein folding environment (Ghosh *et al*, 2019; Dash *et al*, 2025). These were temperature-sensitive (TS) mutants of the *Nourseothricin N-acetyl transferase* (NAT) gene namely TS22, TS15, and TS4, ranked by decreasing severity of misfolding and expressed from a low-copy pRS316 vector. In this study, the same mutants and the wild-type NAT (WtNAT), were genome-integrated into the *yMJ003* derivative strain (Jonikas *et al*, 2009), ensuring a constant genomic copy of the misfolded protein under a constitutive promoter TEF1 (Figure 1A, S1A). The resulting strains were designated YVAM22, YVAM15, YVAM4, and YVAM0, corresponding to the integration of TS22, TS15, TS4, and WtNAT, respectively.

**Figure 1:**
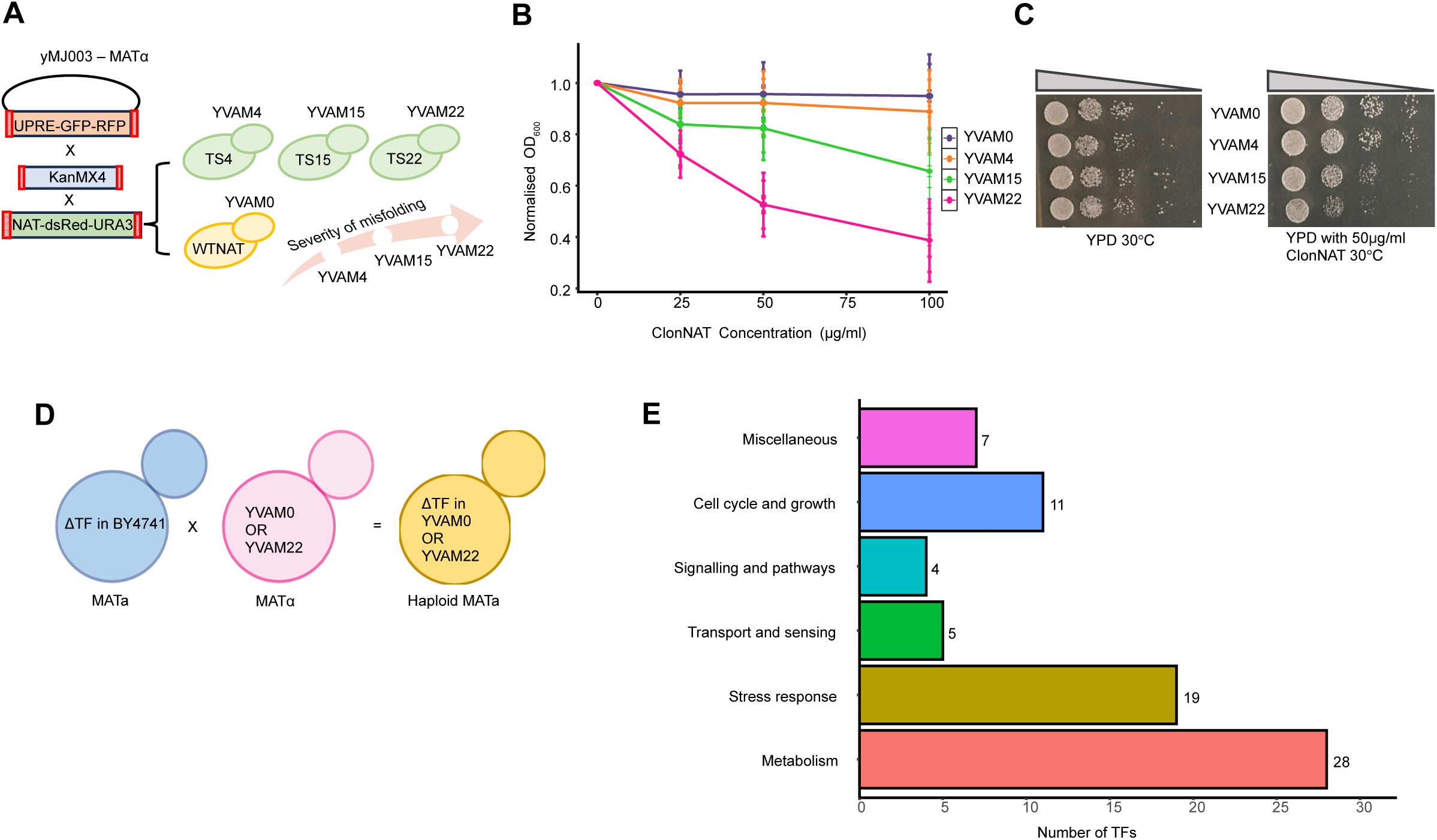
Generation and validation of a genome-integrated protein misfolding sensor for a transcription factor screen. A. Schematic illustrating genome integration strategy: the parental yeast strain (yMJ003 derivative; genotype: MATα, his3Δ1, leu2Δ0, cyh2, met15Δ0, ura3Δ0, LYS+, lyp1Δ::STE3pr-LEU2, ura3Δ::UPRE-GFP-RFP cassette) was sequentially modified. First, the existing UPRE-GFP cassette was replaced with a KanMX cassette, followed by subsequent replacement with the NAT-URA3 cassette containing temperature-sensitive (TS) mutant versions of the NAT gene (TS22, TS15 and TS4). B. Growth of NAT integrated strains was assessed in YPD liquid medium supplemented with increasing concentrations of ClonNAT (25, 50, and 100 µg/ml) at 30°C by optical density (OD) measurements at 600_nm_ after 12-16hrs. Growth was normalized to that in the absence of ClonNAT (n=3). C. Spot dilution assay was performed on YPD agar plates with 50µg/ml ClonNAT conc. and YPD plate as control using 10-fold serial dilutions of each strain grown at 30°C for 36 hours (n=3). D. Schematic representation of genetic cross and selection strategy. MATα haploid YVAM strains carrying genome-integrated NAT mutants were crossed with MATa BY4741 transcription factor deletion strains. Following mating and sporulation, haploid progeny carrying both the NAT mutant and transcription factor deletion markers were selected. E. Functional classification of transcription factors screened in the study. Transcription factors included in the screen were grouped based on their known biological pathways, highlighting the diversity of cellular processes investigated. Categories include metabolism, stress response, transport and sensing, signaling and pathways, cell cycle and growth and miscellaneous functions.

These strains were grown under selection with ClonNAT at 30°C to check for the cells’ ability to tolerate ClonNAT, which in turn is a readout of the cells’ ability to fold the Wt and the mutant NAT proteins; the folding ability is correlated to the proteostasis status of the cell under that condition. All misfolded NAT mutants exhibited reduced growth compared to the wild-type control in both liquid culture (16 hrs) and spot dilution assays (48 hrs), recapitulating prior observations from plasmid-based expression systems (Figure 1B&C) (Ghosh *et al*, 2019). This growth defect was further exacerbated at 37°C, as expected for temperature-sensitive mutants (Figure S1B & S1C). This growth-based readout allowed us to assess how genetic perturbations influence folding capacity. A deletion strain with improved folding of TS22 would exhibit higher growth on ClonNAT while expressing TS22, while impaired folding would result in lower growth. Crucially, the behavior of the WtNAT-expressing strain (YVAM0) must remain consistent across backgrounds to validate that differences are specific to misfolding and not due to confounding effects such as altered expression, translation, or NAT-independent phenomena.

With this model in place, we sought to identify transcriptional regulators involved in maintaining cytosolic protein folding capacity. Using synthetic genetic array (SGA) methodology, we integrated the TS22 sensor and WtNAT into a panel of *S. cerevisiae* transcription factor (TF) deletion strains (Figure 1D). We screened 74 TF deletions, representing diverse cellular functions, including metabolism, stress responses, cell cycle control, growth, epigenetic regulation, transport, and signal transduction (Figure 1E; Figure S1D).

### Folding Defects in Transcription Factor Deletions of Saccharomyces cerevisiae

To evaluate how the deletion of transcription factors (TFs) affects the cellular protein folding environment, we systematically assessed the growth of *Saccharomyces cerevisiae* TF deletion strains expressing either wild-type NAT (WtNAT) or its misfolding-prone variant TS22 under increasing concentrations of ClonNAT (0, 25, and 100 μg/ml) (Figure 2A). Growth kinetics were assessed using two parameters: the maximum growth rate (Mmax) and the time taken to reach this maximum (tMmax) (Figure 2B). To account for inherent differences in growth between strains, we first compared, for each deletion strain and drug concentration, the growth of the TS22 mutant (YVAM22-ΔTF) with that of its corresponding wild-type NAT background (YVAM0-ΔTF). This step provided a strain-specific measure of how the growth rate and timing of the TS22 containing TF deletion strain differed from the wild-type NAT containing strain in the same background. We then normalized these values to the equivalent comparison for the non-deleted control strain (YVAM22/YVAM0), yielding a double-normalized value for each parameter. This approach allowed us to directly assess how much each deletion strain diverged from the non-deleted control in its ability to produce active TS22, independent of baseline growth differences (see Methods for detailed calculation).

**Figure 2:**
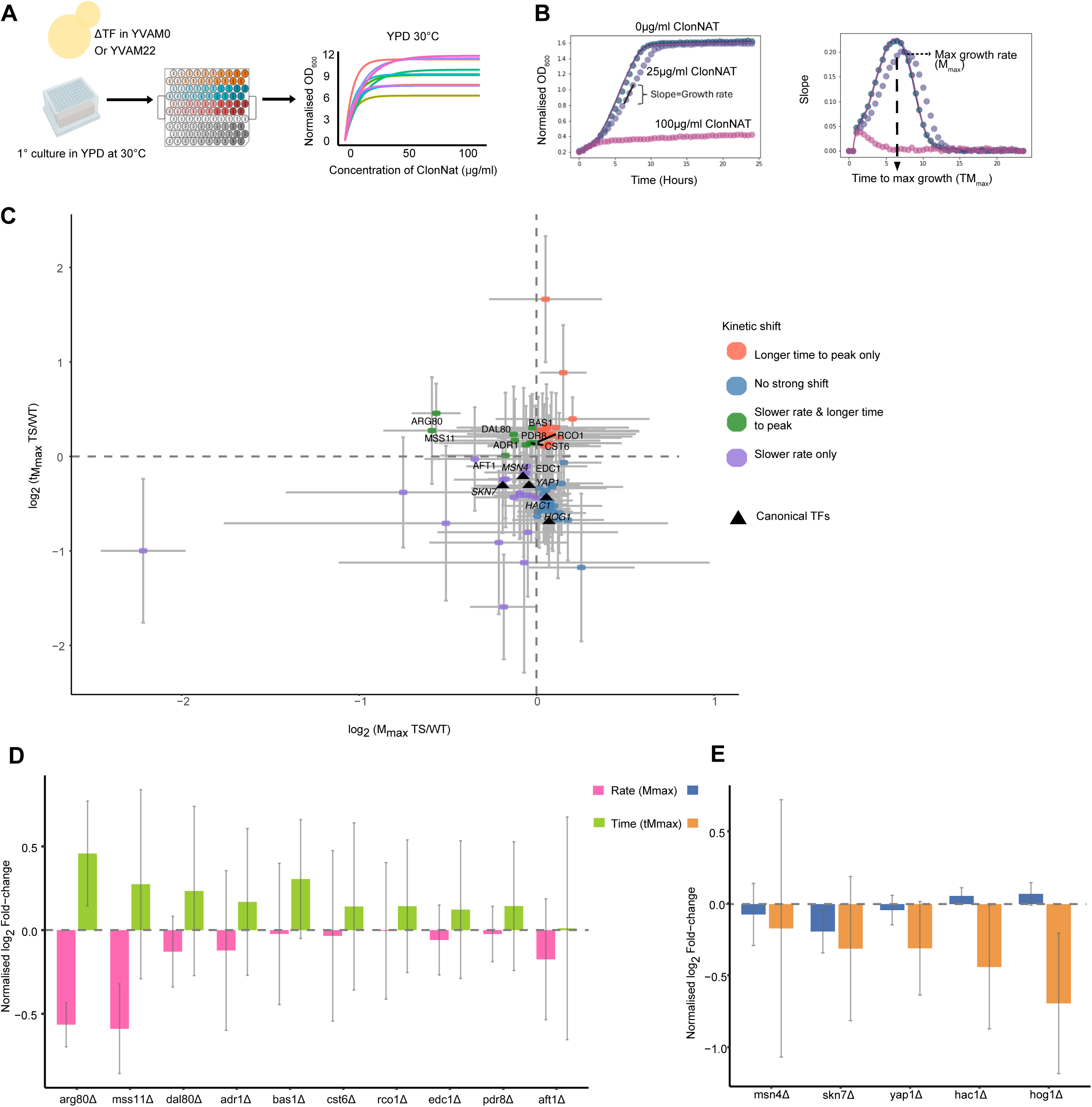
A kinetic growth screen identifies metabolic transcription factors as regulators of proteostasis. A. Schematic representation of the bioscreen assay setup. The setup enabled continuous optical density (OD) measurements at 600nm over 24 hours at 30°C in YPD in different ClonNat concentrations to generate detailed growth curves for YVAM22 and YVAM0 strains with individual transcription factor deletions (n=3). B. Parameters extracted from bioscreen growth curves. Diagram explaining the key growth parameters quantified from bioscreen assays: Maximum growth rate (M_max_): The highest slope of O.D_600nm_ over time; Time to reach maximum growth rate (tM_max_): The time point at which Mmax is achieved. These parameters were used to evaluate the impact of transcription factor deletions on proteostasis in YVAM22 (misfolded protein stress) relative to YVAM0 (wild-type NAT control). C. A two-dimensional log₂–log₂ scatter with dashed reference lines at zero (no change) on both axes. Each dot represents a TF deletion strain. Zero lines divide the plot into four quadrants e.g. the upper-left quadrant (log₂ M_max_ < 0 & log₂ tM_max_ > 0) corresponds to strains exhibiting both slower growth and delayed onset shown as green dots. Orange for deletions which take longer time than wild type to reach maximum growth, blue for deletions which show no strong shift and purple for ones having slower growth only. The black triangle represents canonical TFs screened. Whiskers indicate ± propagated sd values of tM_max_ and propagated standard deviations of M_max_ (see methods). D. Normalised log_2_ Fold change for selected hits from the screen. Green represents growth rate (M_max_) and pink represents time taken to reach maximum growth (tM_max_). E. Normalised log_2_ Fold change for canonical TFs from the screen. Blue represents growth rate (M_max_) and orange represents time taken to reach maximum growth (tM_max_).

Log₂-fold changes derived from these double-normalized values were plotted in a two-dimensional scatterplot (Figure 2C). Dashed reference lines at log₂ = 0 on each axis demarcate no net change and divide the plot into four quadrants, each corresponding to a distinct kinetic-shift phenotype. Strains in the upper-left quadrant (log₂ M□ₐₓ < 0, log₂ t□ₐₓ > 0) exhibited both a reduced growth rate and a delayed onset of growth with respect to YVAM22.

Ten TF deletions prominently clustered in this quadrant: ARG80, MSS11, DAL80, ADR1, BAS1, CST6, RCO1, EDC1, PDR8, and AFT1 (Figure 2C&D, S2A). Notably, these TFs predominantly regulate metabolic or nutrient-responsive pathways. ARG80 is central to arginine metabolism (Dubois *et al*, 1987), MSS11 is involved in carbohydrate utilization (Webber *et al*, 1997), DAL80 modulates nitrogen metabolism (Cunningham & Cooper, 1991), ADR1 regulates ethanol and glucose metabolism (Ciriacy, 1975), and AFT1 controls iron uptake and homeostasis (Yamaguchi-Iwai *et al*, 1995). These results reveal an unexpected, critical dependence on metabolic regulatory networks for maintaining cellular proteostasis and protein folding environment. In contrast, deletions of canonical stress-response transcription factors such as MSN4, SKN7, and YAP1 were predominantly found in the lower-left quadrant, exhibiting a reduced growth rate but an accelerated onset of this impaired growth. This phenotype points to a limited or atypical engagement of classical stress-response pathways under conditions of chronic proteotoxic stress (Figure 2C&E, S2B). TFs such as HAC1 and HOG1, key regulators of the unfolded protein response and osmotic stress, respectively (Cox *et al*, 1997; Rep *et al*, 1999), exhibited minimal shifts (Figure 2C&E), further supporting the notion that TS22-induced misfolding stress preferentially engages alternative regulatory circuits. Together, these findings reveal an unexpected reliance on metabolic transcriptional programs, rather than classical stress signaling pathways, to maintain cytosolic proteostasis. Complete growth parameters and growth curves are provided in Supplementary data (S2A,B).

### Fitness defects upon TF deletions in misfolding stress

To evaluate whether transcription factors identified in the TS22 model also contribute to proteostasis under broader stress conditions, we assessed the growth of selected TF deletion strains in the BY4741 background across diverse proteotoxic environments: heat shock (37°C), tunicamycin-induced ER stress, and azetidine-2-carboxylic acid (AZC), a proline analog that causes widespread protein misfolding (Weids & Grant, 2014). Among the strains tested, only the *arg80Δ* mutant exhibited pronounced growth impairment specifically under AZC treatment (Figure 3A). This suggests that some transcription factors may regulate proteostasis in a stress-specific manner, rather than participating in global folding surveillance. Based on this specificity and consistency, we focused our downstream experiments on Arg80 to better understand its role in proteostasis.

**Figure 3:**
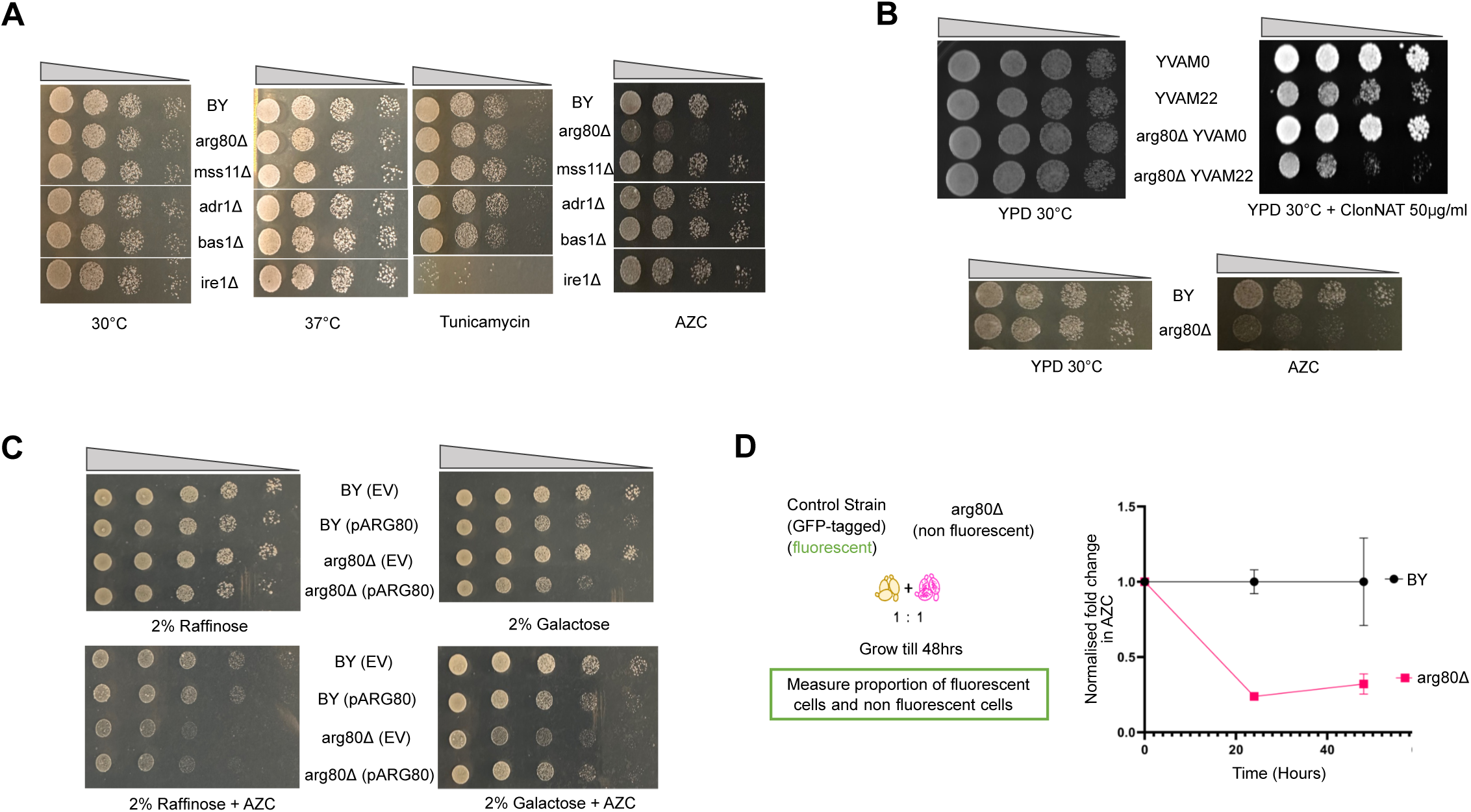
Arg80 is required for cellular fitness under global proteotoxic stress. A. Spot dilution assays of BY4741 and selected transcription factor deletion strains grown at 37°C for heat shock (24 hours), 2.5µg/ml tunicamycin (ER stress) and 4mM AZC (global misfolding stress) with YPD plate at 30°C as control (n=3). B. Spot dilution assays of YVAM and reconstructed arg80Δ in YVAM strains grown in 50 µg/ml ClonNAT (top panel) and BY4741 with reconstructed arg80Δ grown in 2.5 mM AZC (below panel). YPD plate at 30°C was used as a control. Plates were incubated for 36 hours (n=3). C. Spot dilution assay with BY4741 and arg80Δ transformed with Empty vector (EV) and Arg80 Over Expression (pARG80) on synthetic media with 2% Galactose for induction along with 1.5mM AZC. Synthetic media with 2% Raffinose with and without 1.5mM AZC is used as control (n=3). D. Schematic representation of competitive fitness assay in presence of AZC. Competitive fitness assay depicting flow cytometry analysis of BY4741 and arg80Δ populations after 48 hours of growth in YPD media with 4 mM AZC. The relative abundance of GFP+ (BY4741) in black and GFP- (arg80Δ) in pink (normalised fold change) was measured to determine fitness differences under proteotoxic stress(n=3).

To validate that this phenotype was not due to background mutations, we reconstructed the *arg80Δ* allele in both YVAM and BY4741 backgrounds and confirmed the growth defects (Figure 3B). Due to selection compatibility, downstream complementation assays were performed in BY4741 only. Overexpression of ARG80 from a galactose-inducible plasmid restored growth under AZC stress in spot dilution assays, while empty vector controls had no effect (Figure 3C, S3A). We further quantified this fitness defect using a competitive fluorescence-based assay. GFP-tagged wild-type cells (tagged at the TDH2 locus, TDH2-GFP) were co-cultured with untagged *arg80Δ* in a 1:1 ratio and grown in the presence or absence of AZC. Tracking the populations with flow cytometry over 48 hours, we observed that *arg80Δ* cells were steadily outcompeted by GFP-tagged wild type cells (Figure 3D), specifically only in the presence of AZC, confirming the reduced fitness of *arg80Δ* under proteotoxic stress. Together, these results establish Arg80 as a central regulator of proteostasis, acting across both localized (cytosolic folding of TS22) and global protein misfolding contexts.

### Arg80 is regulated by misfolding stress

While we find some of the TFs to play a role in maintaining proteostasis through the TS22 activity assay, it is not imperative that these TFs are activated during misfolding stress. A bona fide proteostasis-regulating TF would also have the potential to be regulated during conditions that perturb proteostasis. To assess whether the transcription factors (TFs) identified through our screen are functionally responsive to misfolded protein stress, we examined their regulatory activity in RNA-seq data from cells experiencing misfolding stress. Gene expression profiles of yeast strains, each expressing one of the three NAT misfolding mutants (YVAM22, YVAM15, and YVAM4) were compared against the wild-type control (YVAM0) to evaluate whether known targets of each TF exhibited transcriptional changes consistent with their established regulatory roles. A schematic overview of the analysis is shown in (Figure 4A). For each TF, using a log_2_ fold change cutoff of |0|, we calculated the percentage of activated targets that were upregulated and repressed targets that were downregulated under stress conditions. The average of these two percentages provided a composite score named as TF activity score. (Figure 4B). Additionally, a Wilcoxon rank-sum test was used to statistically compare the log_2_ fold changes of each TF’s targets with those of other genes (Figure 4C). Taken together, these reflect the extent and direction of TF’s regulatory activity.

**Figure 4:**
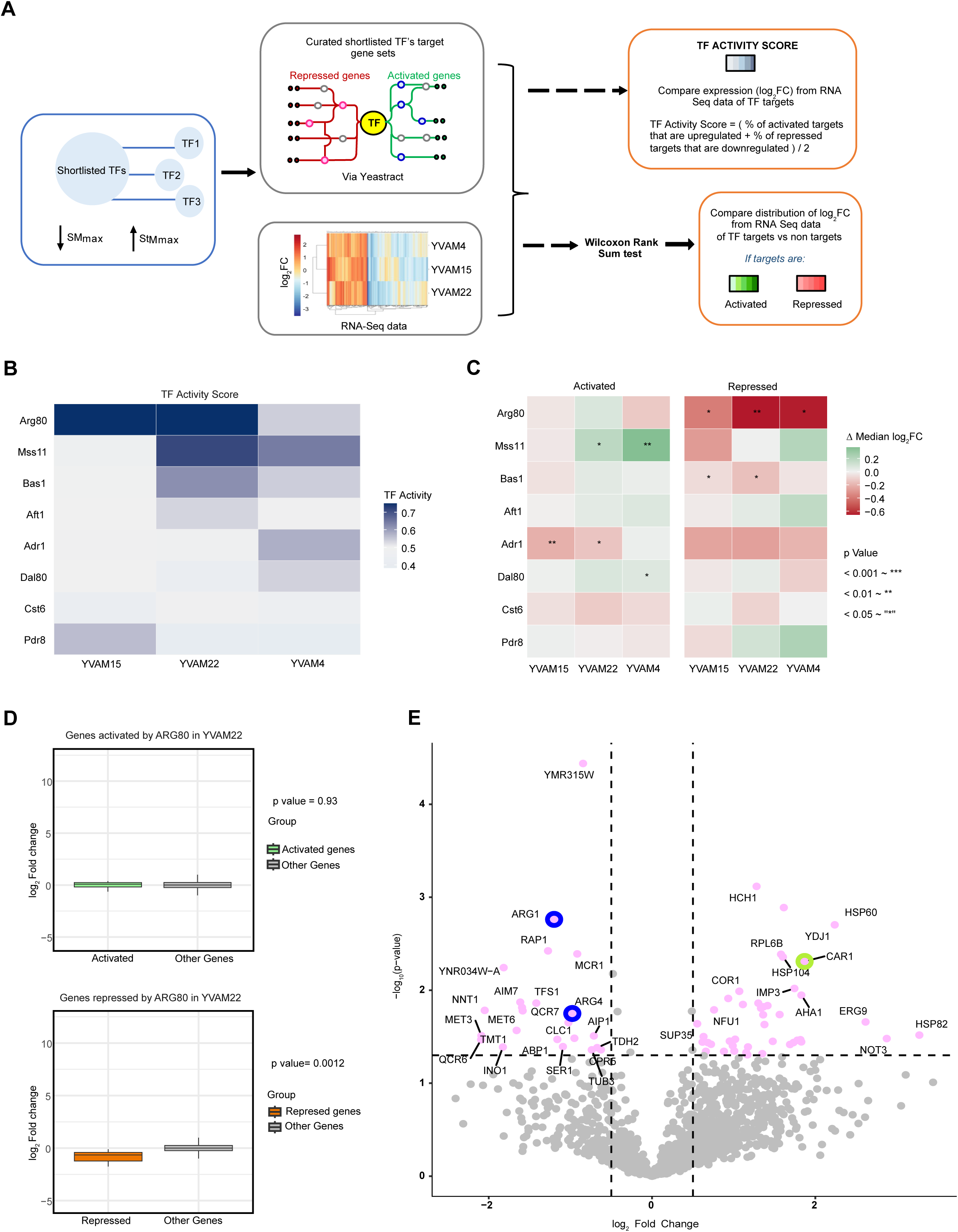
Misfolding stress activates Arg80. A. Schematic of TF activity analysis combining bioscreen and RNA-seq data. B. TF activity score heatmap for selected TFs. Darker shade of blue represents higher TF activity in presence of NAT mutants. C. TF activity separated by activation and repression of genes by TF in presence of NAT mutants. D. Box plots show the distribution of log_2_ fold change (YVAM22 vs. YVAM0) for genes activated by Arg80, genes repressed by Arg80, and all other genes (Rest). Green are activated genes, orange represent repressed genes and grey shows rest (other) genes. The central line indicates the median. p-values were determined using the Wilcoxon rank-sum test. E. Proteins significantly altered upon AZC treatment in BY4741 are labeled in the volcano plot (log₂FC > 0.5 or < −0.5, −log₁₀ p-value > 1.3). Arginine biosynthesis proteins (repressed by Arg80) are highlighted with a blue circle, while arginine catabolism proteins (activated by Arg80) are highlighted with a green circle (n=3).

Among the TFs analyzed, Arg80 exhibited consistent regulatory behavior across all three mutants, marked by the significant downregulation of its repressed targets (Figure 4C&D, &S4A). This pattern indicates that Arg80 remains active or is further induced under misfolded protein stress. Mss11 showed activation in YVAM22 and YVAM4, while Bas1 was active in YVAM15 and YVAM22 (Figure 4C and S4B). Other TFs displayed weaker trends or effects contrary to their canonical roles (Figure 4C). Arg80 is known to regulate arginine metabolism, repressing biosynthetic genes (ARG1, ARG3, ARG5,6, and ARG8) and inducing catabolic genes (CAR1 and CAR2) in response to arginine (Dubois *et al*, 1987). Given its consistent activation in all NAT misfolding mutants, we investigated whether Arg80 was also activated under other proteotoxic stress. In BY4741, treatment with azetidine-2-carboxylic acid (AZC) led to upregulation of arginine catabolic proteins and suppression of biosynthetic proteins, as shown by quantitative proteomics (Figure 4E). These results indicate that Arg80-mediated regulation is a general feature of the cellular response to misfolded protein stress and may represent a parallel adaptive pathway in proteostasis regulation.

### Arginine as one of the effectors during activation of Arg80 in misfolded protein stress

Arg80 functions as a transcription factor that regulates arginine metabolism by repressing biosynthesis and activating catabolism, making arginine a strong candidate as an effector in misfolded protein stress. Consistent with this, we observed a significant increase in intracellular arginine levels in *arg80Δ* under AZC stress, as well as its accumulation in YVAM22 upon Arg80 deletion, suggesting that Arg80 loss disrupts arginine homeostasis (Figure 5A). To assess whether this accumulation affects proteostasis, we supplemented exogenous arginine in growth assays and observed that as arginine concentration increased, the growth of YVAM22 worsened in ClonNAT assays, with no effect on YVAM0; thus activity of TS22 decreased in vivo without affecting WtNAT, indicating that arginine levels compromises folding of folding-compromised protein in the cytosol (Figure 5B). A similar trend was observed in YVAM15, a moderate misfolding mutant, which exhibited reduced growth upon 1mM arginine supplementation, whereas YVAM4, a mild misfolding mutant, remained unaffected (Figure S3C). These findings suggest that the impact of arginine accumulation on proteostasis may be dependent on the severity of the folding defect of the misfolding mutants of the same protein. ARG80 induces the expression of CAR1 to metabolise arginine. If the proteostasis defect in *arg80Δ* is due to the cell’s inability to catabolize arginine, CAR1 deletion should mimic arg80 deletion. Deletion of CAR1, which encodes arginase and converts arginine to ornithine, resulted in phenotypic defects similar to *arg80Δ*. In the YVAM background, CAR1 deletion decreased the activity of the misfolding mutant TS22 without altering the activity of WtNAT (Figure 5C). Extending this observation, we examined CAR1’s role in a global misfolding stress model by deleting CAR1 in BY4741 and subjecting cells to AZC stress. Similar to *arg80Δ*, *car1Δ* exhibited heightened sensitivity to AZC and an accumulation of arginine (Figure 5D.i&ii), reinforcing the link between arginine metabolism and proteostasis regulation. Interestingly, targeted metabolomics revealed that ornithine levels were significantly depleted in both *arg80Δ* and *car1Δ* strains under AZC stress, raising the possibility that ornithine deficiency contributes to proteostasis defects (Figure 5E.i). However, despite its depletion, supplementing ornithine did not rescue the growth defects of *arg80Δ* or *car1Δ*; instead, it further exacerbated growth impairment under AZC conditions (Figure 5E.ii). This suggests that the observed phenotype is not simply due to a lack of ornithine but rather due to accumulation of arginine. Finally, when we overexpressed Car1 in *arg80Δ*, restoring arginine catabolism, we observed a significant rescue of the growth phenotype in AZC stress (Figure 5F & S3B), confirming that metabolic dysregulation, and catabolism of arginine plays a pivotal role in the proteostasis defects observed in *arg80Δ*. Together, these findings establish Arg80 as a key modulator of proteostasis via metabolic regulation, demonstrating that amino acid homeostasis is tightly linked to the cell’s ability to manage misfolded protein stress.

**Figure 5:**
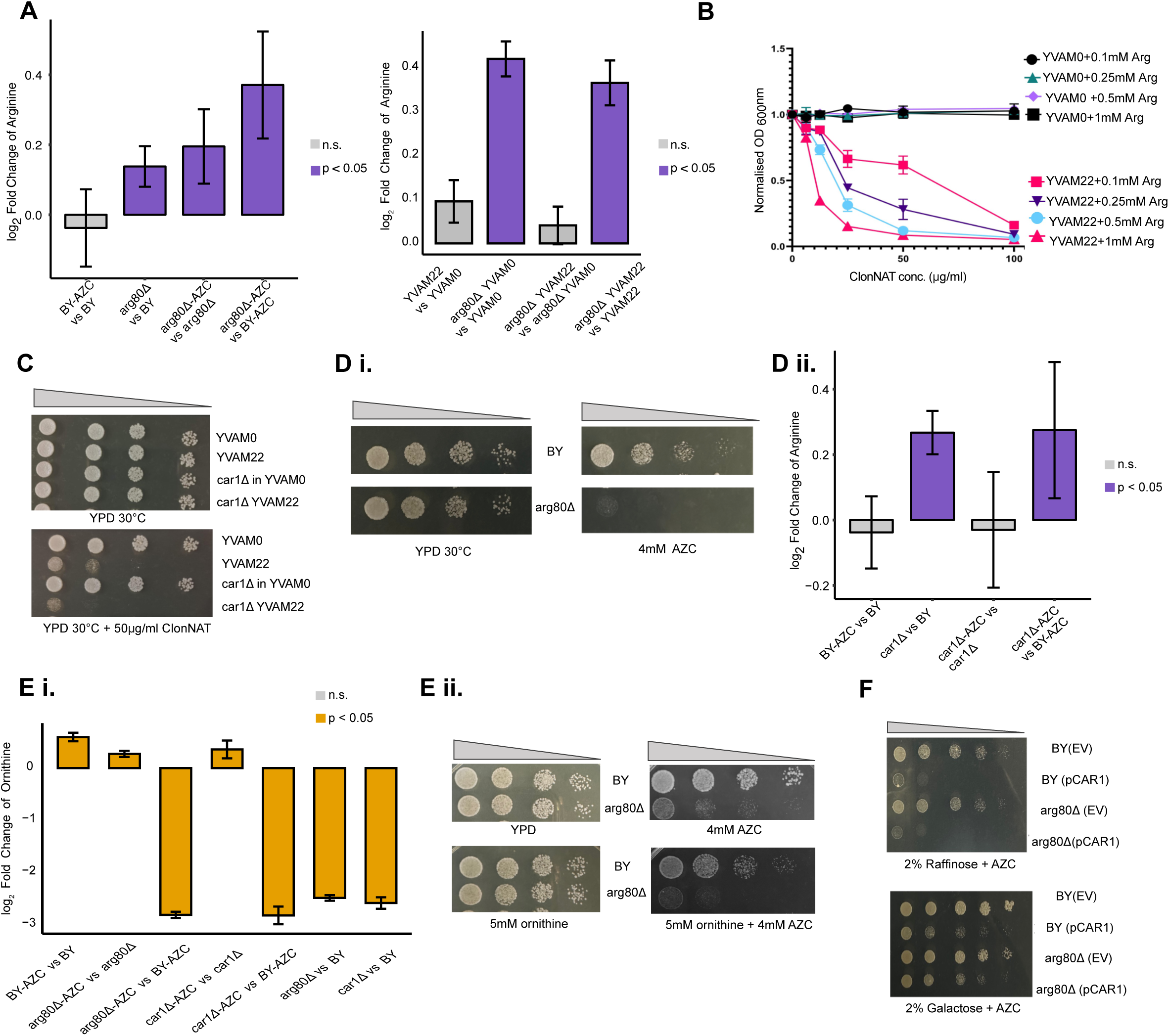
Arginine accumulation compromises proteostasis in arg80Δ mutants. A. Targeted metabolomics analysis of arginine levels in BY4741 and arg80Δ with AZC treatment (left panel) and in arg80Δ in YVAM22 vs YVAM22 (right). The y-axis represents log₂ fold change, while the x-axis shows individual metabolite levels. Bars highlighted in purple indicate significant changes (p < 0.05, determined using a two-sample Welch’s t-test on log₂-transformed replicates). B. ClonNAT assay of YVAM0 and YVAM22 grown in increasing concentrations of arginine (0.1mM – 1mM) under ClonNAT selection (0, 12.5, 25, 50, and 100 µg/ml) at 30°C in complete synthetic media. C. Spot dilution assay of YVAM0, YVAM22, and car1Δ in these strains on 50 µg/ml ClonNAT in YPD at 30°C. YPD plates at 30°C were used as a control. Plates were incubated for 36 hours. D. Spot dilution assay of BY4741 and car1Δ with 4mM AZC. YPD plates at 30°C were used as a control. Plates were incubated for 36 hours (i). Targeted metabolomics analysis of arginine levels in BY4741 and car1Δ with AZC treatment. The y-axis represents log₂ fold change, while the x-axis shows individual metabolite levels. Bars highlighted in purple indicate significant changes (p < 0.05, determined using a two-sample Welch’s t-test on log₂-transformed replicates)(ii). E. (i) Targeted metabolomics analysis of ornithine levels across strains with AZC treatment. The y-axis represents log₂ fold change, while the x-axis shows individual metabolite levels. Bars highlighted in yellow indicate significant changes (p < 0.05, determined using a two-sample Welch’s t-test on log₂-transformed replicates).(ii) Spot dilution assay in 4mM AZC with 5mM ornithine supplementation. Controls included only ornithine, only AZC, and YPD at 30°C. Plates were incubated for 36 hours. F. Spot dilution assay with BY4741 and arg80Δ transformed with Empty vector (EV) and CAR1 Over Expression (pCAR1) on synthetic media with 2% Gal (SG) for induction along with 1.5mM AZC. Synthetic media with 2% Raffinose (SR) with and without 1.5mM AZC is used as control.

### Arg80 and Gcn2 Mediate a Novel Proteostasis Axis Through Metabolic and Translational Regulation

To deepen our understanding of Arg80’s role as a modulator of proteostasis, we examined proteome changes in YVAM22 following the deletion of arg80. Given our previous findings indicating arginine as a potential effector molecule contributing to the compromised activity of misfolded proteins in *arg80Δ*, we sought to elucidate the downstream pathways affected by its absence. By investigating other altered pathways, we aimed to gain insights into any additional roles of Arg80. As expected, upon deletion of arg80 we saw an upregulation of proteins related to arginine biosynthesis like ARG1, ARG3, ARG5 and ARG8 and downregulation of proteins related to arginine catalysis like CAR1 (Figure 6A). Additionally, we observed enrichment of proteins in the starch and sucrose metabolism category (Figure 6B). These proteins were mainly involved in trehalose biosynthesis, like TPS1, TPS2, TPS3, etc. The upregulation of trehalose biosynthesis was unexpected in the context of arginine metabolism, as there is no known interaction between arginine and trehalose in the proteostasis network. However, separate evidence suggests that both arginine and trehalose can function as small molecules or chemical chaperones (Tapia & Koshland, 2014; Baynes *et al*, 2005). Consistent with this, AZC treatment of *arg80Δ* and *car1Δ* strains resulted in a significant increase in trehalose levels compared to BY4741 (Figure 6C). To explore the relationship between Arg80 and trehalose, we depleted trehalose biosynthesis genes in the *arg80Δ* strain. Interestingly, the double deletion of ARG80 and TPS2 restored growth to levels comparable to the parental BY4741 strain under AZC stress, whereas single deletions exacerbated the growth defect (Figure 6D). These results reveal an unexpected connection between ARG80 and trehalose biosynthesis, suggesting that Arg80’s role in proteostasis extends beyond arginine metabolism. The rescue of growth defects in the *arg80Δtps2Δ* strain under AZC stress points to a potential compensatory mechanism between trehalose biosynthesis and arginine regulation in maintaining cellular homeostasis.

**Figure 6:**
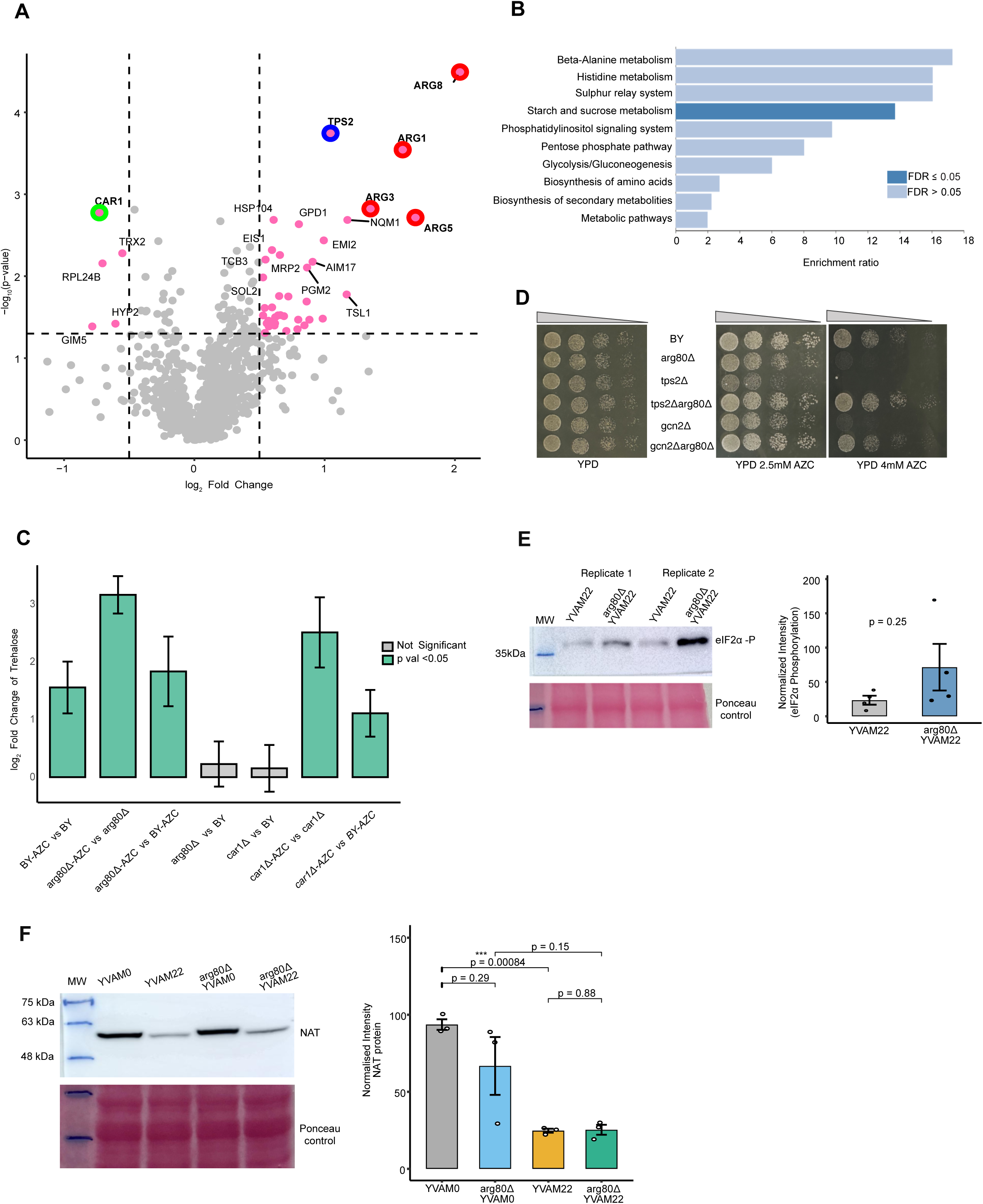
Arg80 connects arginine metabolism to trehalose biosynthesis and the Gcn2-mediated stress response. A. Volcano plot depicting significantly altered proteins in arg80Δ in YVAM22. Labels indicate significant proteins (log₂FC > 0.5 or < −0.5, −log₁₀ p-values > 1.3). Arginine biosynthesis related proteins are upregulated circled in red. Arginine catabolic genes downregulated and circled in green, trehalose biosynthesis proteins circled in blue. B. Over-representation analysis of significantly altered proteins using www.webgestalt.org. The bar graph highlights the enrichment (FDR≤0.05) of starch and sucrose metabolism pathways, including proteins primarily involved in trehalose biosynthesis. C. Targeted metabolomics analysis of trehalose levels in BY4741, arg80Δ, and car1Δ under AZC stress. The y-axis represents log₂ fold change, while the x-axis shows individual metabolite levels. Bars highlighted in green indicate significant changes (p < 0.05, determined using a two-sample Welch’s t-test on log₂-transformed replicates) D. Spot dilution assay of BY4741, arg80Δ, tps2Δ, arg80Δtps2Δ, gcn2Δ, arg80Δgcn2Δ grown on YPD agar plates supplemented with 2.5mM and 4mM AZC at 30°C. YPD agar plate at 30°C served as controls. Plates were incubated for 36 hours. E. Western blot analysis of eIF2α expression in arg80Δ in YVAM22 and YVAM22. Ponceau staining was used as a loading control (left panel). The right panel shows quantification of the western blot signal using ImageJ (*n* = 4). F. Western blot analysis of NAT expression in arg80Δ in YVAM22 and YVAM22 at 0.6 O.D_600_ Ponceau staining was used as a loading control (left panel). The right panel shows quantification of the western blot signal using ImageJ (*n* = 3).

Given the observed metabolic alterations and the established role of the nutrient-sensing kinase Gcn2 in regulating translation during amino acid imbalance, we explored potential cross-talk between ARG80-mediated metabolism and Gcn2-dependent translational regulation. Indeed, the double deletion of ARG80 and GCN2 also ameliorated the fitness defects under AZC-induced proteotoxic stress (Figure 6D). Supporting a translational stress connection, Western blotting revealed increased phosphorylation of eIF2α in *arg80Δ* in YVAM22 compared to the parental strain (Figure 6E). However, to assess whether this led to any direct translation defects, we examined steady-state levels of NAT in *arg80Δ* in YVAM22 and compared them to YVAM22. The unchanged NAT protein levels between these strains (Figure 6F) suggest that eIF2α phosphorylation does not result in global translational repression, at least for NAT. Instead, the proteotoxicity in *arg80Δ* appears to arise from metabolic imbalance rather than impaired translation of the misfolded protein itself. The rescue observed in *gcn2Δarg80Δ* suggests that Gcn2-mediated stress responses, rather than direct translational inhibition, might contribute to the proteostasis defects in *arg80Δ*. Specifically, deletion of ARG80 alters arginine homeostasis, potentially increasing AZC incorporation or disrupting metabolic pathways linked to proteotoxicity. Concurrently, deletion of GCN2 could alleviate amino acid sensing–dependent stress responses, thereby buffering the effects of altered arginine metabolism. These findings highlight how metabolic and translational pathways converge to influence proteostasis, revealing Arg80 as an unconventional player in proteostasis regulation beyond canonical chaperone and degradation networks.

## DISCUSSION

Proteostasis is a fundamental cellular process that ensures proteins fold correctly and maintain functionality. While traditionally viewed as a network governed by molecular chaperones and degradation systems, growing evidence suggests that metabolic pathways also play a critical role in maintaining proteostasis. However, the mechanisms by which metabolic regulators influence protein homeostasis remain largely unexplored. In this study, we identify Arg80, a transcription factor primarily known for its role in arginine metabolism, as an unexpected regulator of proteostasis in *Saccharomyces cerevisiae*. Our findings reveal that Arg80’s function extends beyond amino acid regulation, linking metabolic balance to cellular responses to misfolded protein stress.

Under normal conditions, Arg80 regulates arginine biosynthesis and catabolism to maintain amino acid homeostasis. However, intracellular arginine levels increase significantly in its absence, leading to metabolic imbalances that sensitize cells to misfolded protein stress. The phenotypic similarity between *arg80Δ* and *car1Δ* mutants, where CAR1 encodes arginase (Sumrada & Cooper, 1984), the enzyme responsible for converting arginine to ornithine, suggests that proteostasis defects stem from defective arginine catabolism. The failure of exogenous ornithine supplementation to rescue *arg80Δ* defects further supports the notion that controlled accumulation of arginine is crucial for cellular homeostasis.

Our study also uncovers an unexpected link between arginine metabolism and carbohydrate stress responses; both *arg80Δ* and *car1Δ* mutants exhibit increased trehalose biosynthesis under misfolded protein stress. Trehalose, a well-established chemical chaperone, protects proteins from aggregation in various stress conditions. However, in *arg80Δ* mutants, trehalose accumulation correlates with exacerbated stress sensitivity rather than protection. The genetic rescue observed upon deletion of *TPS2*, a key enzyme in trehalose biosynthesis (Bell *et al*, 1992; De Virgilio *et al*, 1993), suggests that trehalose overproduction is not inherently protective but rather a maladaptive stress response in the presence of excess arginine, triggered by metabolic imbalance. These findings highlight a broader, underappreciated crosstalk between amino acid metabolism and carbohydrate stress responses, reinforcing the idea that proteostasis extends beyond classical chaperone networks and is dynamically regulated by metabolic state.

Beyond metabolic regulation, our study reveals a functional interaction between Arg80 and Gcn2, a well-characterized sensor of amino acid imbalance and translational stress (Wek *et al*, 1995; Deng *et al*, 2002). Loss of Arg80 led to increased phosphorylation of eIF2α, a hallmark of Gcn2 activation, despite the paradoxical presence of elevated intracellular arginine. One possible explanation is that dysregulated arginine metabolism disrupts proper tRNA charging, leading to a starvation-like signal that activates Gcn2. This aligns with previous findings showing that amino acid imbalances, not just deprivation, can induce translational stress responses (Krokowski *et al*, 2022; Wek *et al*, 2006; Harding *et al*, 2003). The suppression of *arg80Δ* phenotypes upon *GCN2* deletion confirms that aberrant activation of the integrated stress response contributes to proteostasis defects (Costa-Mattioli & Walter, 2020). However, steady-state levels of the misfolded NAT protein remained unchanged in *arg80Δ*, suggesting that eIF2α phosphorylation does not globally inhibit translation but may selectively alter the synthesis of stress-responsive proteins. This highlights an unexpected metabolic-translational axis in proteostasis regulation.

These findings have significant implications beyond yeast. In higher eukaryotes, amino acid metabolism and translational control mechanisms are tightly linked to neurodegenerative diseases, aging, and cellular stress responses (Wiebe *et al*, 2020; Ghosh *et al*, 2020; Ling *et al*, 2023). The Gcn2-eIF2α pathway, conserved in mammals, has been implicated in disorders such as Alzheimer’s, Huntington’s, and Parkinson’s, where chronic activation contributes to neuronal dysfunction (Ma *et al*, 2013; Bravo-Jimenez *et al*, 2025; Costa-Mattioli & Walter, 2020). Our results suggest that misregulated amino acid metabolism could influence proteostasis decline in human cells through mechanisms similar to those observed in yeast. Additionally, trehalose, which accumulates in *arg80Δ* mutants, has been widely studied for its neuroprotective properties (Sevriev *et al*, 2024; Yap *et al*, 2023). While trehalose supplementation has been shown to reduce protein aggregation and enhance autophagy in neurodegenerative models (Sarkar *et al*, 2007), our findings suggest that its endogenous accumulation may not always be beneficial and could represent a stress-induced compensatory response rather than a direct protective mechanism. This highlights the importance of understanding the cellular context in which trehalose contributes to proteostasis.

Despite its insights, this study has certain limitations. Our genetic screen focused on a subset of transcription factors, leaving open the possibility of additional, undiscovered proteostasis regulators. Moreover, single deletion screens do not capture epistatic interactions, meaning that combinatorial regulatory effects remain to be fully explored. Further studies are needed to clarify the precise molecular mechanisms by which Arg80 interfaces with metabolic and stress response pathways, particularly its influence on translation and proteostasis beyond the Gcn2 axis.

Several open questions arise from this work. Does trehalose accumulation in *arg80Δ* mutants actively exacerbate stress, or is it simply a byproduct of metabolic dysregulation? Is there a problem in folding kinetics in the presence of excess arginine, that is exacerbated in the presence of excess trehalose? Furthermore, extending these findings to mammalian systems could provide valuable insights into how amino acid metabolism influences proteostasis in higher eukaryotes. If metabolic imbalances similar to those in *arg80Δ* mutants contribute to proteostasis decline in human cells, interventions targeting amino acid homeostasis or translational stress pathways could hold therapeutic potential. It is of importance to note that ARG1, the human homolog of yeast CAR1, is implicated in hyperargininemia (Haraguchi *et al*, 1990; Grody *et al*, 1992) is linked to spastic paraparesis, progressive impairment of intellectual ability, and the nervous system. If this results from a proteostasis insult, as many neurodegenerative diseases do, would need to be investigated in light of our findings.

In conclusion, our study establishes Arg80 as a key integrator of metabolic and translational responses in proteostasis regulation. By coordinating amino acid metabolism with stress adaptation mechanisms, Arg80 ensures that biosynthetic processes remain aligned with protein quality control. The discovery of its role in modulating the Gcn2-eIF2α pathway and trehalose biosynthesis expands our understanding of how metabolic state influences proteostasis beyond canonical chaperone networks. More broadly, our findings suggest that metabolic interventions could be harnessed to modulate proteostasis capacity in both yeast and higher eukaryotes, offering new avenues for targeting metabolic pathways in protein misfolding diseases.

## METHODS

### Growth Measurements and Data Analysis

From an overnight primary culture, cells were inoculated into 96-well deep-well plates containing 400 µL of medium per well at an initial optical density (OD₆₀₀) of 0.05. Cultures were grown at 30 °C with shaking at 200 rpm for 12hrs or 16hrs in the presence of varying concentrations of the antibiotic ClonNAT. Growth was monitored using a multi-plate reader (TECAN). Assays with L-azetidine-2-carboxylic acid (AZC) were performed in the same manner. For the genetic screening using the Bioscreen C system, 200 µL cultures containing varying concentrations of the relevant chemical (antibiotic, AZC, or other treatment) were grown at 30°C or 37°C in honeycomb plates, with shaking at normal amplitude. Absorbance at 600_nm_ was recorded at 30- or 60-minute intervals for 24hrs using a Thermo Labsystems Type FP-1100-C Bioscreen C Automated Microbiology Growth Curve Analysis System.

### Growth parameter extraction

Growth curves for each strain were obtained from three independent biological replicates (*n* = 3). For each replicate, two parameters were extracted:

1. M_max_ – the maximum observed growth rate.
2. tM_max_ – the time taken to reach Mmax.

#### Double normalization procedure

To quantify strain-specific effects attributable to the TS22 mutant, we used a two-step normalization approach.

- **Step 1 – Within**-**background normalization:** For each deletion strain and drug concentration, the growth parameter of the TS22 mutant (YVAM22) was divided by the corresponding value from its wild-type NAT background (YVAM0). This yielded:

- The *M_max_ ratio* – M_max_(TS deletion strain) ÷ M_max_(WT deletion strain)
- The *tM_max_ ratio* – tM_max_(TS deletion strain) ÷ tM_max_(WT deletion strain)
- **Step 2 – Control**-**strain normalization:** These ratios were then divided by the corresponding ratios obtained for the non-deleted control strain (YVAM). This “double normalization” controlled for baseline strain-specific growth differences and provided a direct comparison of each deletion strain to the non-deleted control in its ability to produce active TS22.

Double-normalized M_max_ = [M_max_(TSdel) / M_max_(WTdel)] ÷ [M_max_(TSyvam) / M_max_(WTyvam)]
Double-normalized tM_max_ = [tM_max_(TSdel) / tM_max_(WTdel)] ÷ [tM_max_(TSyvam) / tM_max_(WTyvam)]

#### Propagation of uncertainties for fold-change calculations

Standard deviations (SD) for fold-changes were calculated from biological replicates using the delta method. Specifically, for a fold-change (FC) defined as the ratio of the mean value for TS22 to the mean value for the corresponding wild-type (WT), the SD was estimated as:

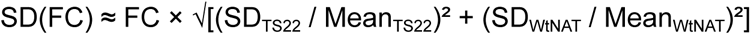

where *Mean* and *SD* refer to the mean and standard deviation, respectively, of the biological replicate measurements for each strain.

### Western blotting

Secondary culture was inoculated at ∼ 0.2 O.D. and grown at 30 °C; 200 rpm. After 6 h of growth, the cultures were pelleted. Protein was isolated by alkaline lysis method, concentration was estimated by BCA (Thermo Scientific, USA) and 30 μg of total protein was loaded per lane in a 10% gel. The nitrocellulose/ PVDF membrane was probed with a custom anti-Nat-r primary antibody (1:5000 dilution). In case of GFP, primary antibody (1:10,000 dilution). A horseradish peroxidase-conjugated goat anti-rabbit antibody was used as a secondary antibody and developed through chemiluminescence (Immobilon Western Chemiluminescent HRP Substrate, Merck Millipore). Blots were quantitated using ImageJ and were normalized to ponceau with its respective blot. For cycloheximide (CHX) chase, cells were inoculated at ∼ 0.2 O.D. and grown at 30°C; 200 rpm in secondary culture. After 4 h of growth, cycloheximide (0210018305, MP Biomedicals) was added to a final concentration of 40 ug/ml and aliquots collected (kept in −20°C) and mentioned time intervals. These cultures were then pelleted, protein isolated and processed for western blotting.

### RNA isolation

Secondary culture was inoculated at 0.2 O.D600nm in 50 ml of media from overnight grown primary culture and was allowed to grow till 0.6 O.D600nm. Cultures were pelleted down at 4°C at 4000 rpm and washed with chilled 1xPBS and then transferred to MCT. Autoclaved acid washed beads were added twice the amount of pellet with 1 ml Trizol and resuspended. Bead beating for 10 cycles with 1 min OFF on ice and 1 min ON. After centrifugation at 8000 rpm, supernatant was collected in fresh MCT. To this 800μl trizol lysate, 250μl chloroform was added and vortexed. The contents should turn into rose milk color only then should proceed further. The samples were then allowed to incubate for 10 min at RT. After centrifugation at 13000 rpm at 4°C supernatant was collected without touching the middle layer. Equal volumes of chilled 85% ethanol was added to the sup and mixed gently by pipetting. The contents were then transferred to the qiagen column from RNeasy Mini Kit and spun at 10000 rpm for 30s and then proceeded as per kit protocol. Note* Don’t store cell pellets, proceed on the same day. Always use freshly double autoclaved tips and MCTs. Before loading on gel heat equal amount of RNA and loading buffer at 70°C for 2-3 min and snap chill on ice for 5 min. Store RNA at −80. If isolated for qRT PCR, proceed with cDNA preparation on the same day.

### RNA Sequencing

For RNA SEQ, first mRNA was purified from cultures using DynabeadsTM mRNA Purification Kit. Protocol was the same as RNA isolation till addition of glass beads. Instead of Trizol, lysis buffer provided in the kit was added (300μl for 50ml culture and 500μl for 100ml culture). Bead beating for 5 cycles 1min OFF on ice and 1min ON. Centrifuged at 10000 rpm for 2min, collect sup. 200μl of beads (1mg) will bind to 2μg of mRNA, so 60μl of beads were taken for each sample MCT. Beads were washed and equilibrated with the lysis buffer by placing them on a magnetic stand and then lysate sup was added and incubated at RT for 15 min in rotation. After incubation MCT was again placed on a magnetic stand and washed using wash buffers A and B provided in kit 2 times each (500μl volume). For elution 20μl of elution buffer (10mM Tris-HCl) was used. The samples were heated at 70°C after addition of the elution buffer for 2 min and then placed on a magnetic stand and sup was collected and quantified using nanodrop. Library for RNA sequencing was then prepared following Ion Total RNA-Seq Kit v2 USER GUIDE and run on Ion Proton sequencer.

### Data Availability

The RNA-seq data generated in this study have been deposited in the NCBI BioProject database under accession number PRJNA1144243.

### Targeted metabolomics

Primary overnight culture from fresh YPD plate was used to set up secondary culture with 0.2 as initial O.D600nm and allowed to grow till exponential phase (∼0.6 O.D600nm). Cells were harvested at 4°C and pellets were washed with LS/MS grade water. Cell pellets were resuspended in 500*μ*l pre-chilled 80% methanol. The mixture was allowed to snap freeze and thawed using liquid nitrogen. Acid washed glass beads were added the same amount as the size of the pellet and bead beating was done for 10 cycles with 1 min ON and 1min OFF on ice. Centrifuged at 10,000 rpm at 4°C for 15 min and sup was transferred to fresh MCT. The sample was then subjected to vacuum drying at V-AQ at 30°C till the pellet is completely dry (usually takes 4-5 hours). The pellet here can be stored at −80°C but it is preferred to run the samples as soon as possible. We always ran the samples the next day. Samples were brought from −80°C in ice and resuspended in 250-300*μ*l and vortexed. The samples were then centrifuged at 10000 rpm for 10 min and 35*μ*l was taken in mass spec compatible vials. Note *For each sample 5 biological replicates were taken. For a single run the same media was used. All the microtips and MCTS used were unautoclaved. Samples were run on TSQ AltisTM Triple Quadrupole Mass Spectrometer (Cat no. TSQ02-10002) using ACQUITY UPLC BEH HILIC Column, 130Å, 1.7 μm, 2.1 mm X 50 mm, 3/pk (Cat no. 176001092).

### Competitive Fitness Assay

For competitive fitness assay, three single colonies of each strain were inoculated in 400*μ*l of YPD broth in a deep well plate (having a well volume of 2 ml each), along with Tdh2-GFP (unevolved fluorescent control where Tdh2 is GFP tagged). The cultures were grown overnight at 30°C on 200 rpm as primary cultures. The assay was set up by mixing an equal number of cells of both the fluorescent (Tdh2-GFP) and untagged strain in a well, as monitored by their O.D600nm i.e., 0.4 O.D600nm was set for both the strains and 100*μ*l of each was mixed with 200*μ*l of YPD media. The cultures were grown and passaged after every 12 hours. The proportion of each population was measured by Flow cytometer BD LSR II after every 24 hours till 72 hours (or as specified). For assessing the fitness of the cells in AZC, 2.5mM or 4mM of AZC concentration was used throughout the study. The cultures were grown at 30°C in an incubator at 200 rpm.

## ACKNOWLEDGEMENTS

We thank Dr. Bani Jolly for helpful discussions and initial assistance with RNA-seq data analysis and Dr. Praveen Singh for assistance with the proteomics run and its analysis. K.C. acknowledges funding from CSIR projects OLP2303 and OLP2504. We acknowledge CSIR-IGIB for infrastructural support. Fellowship support is gratefully acknowledged from the UGC (A.S., S.R.), CSIR-IGIB (Z.Z.), and CSIR (K.P., V.S.).

## AUTHOR CONTRIBUTIONS

A.S. contributed to conceptualization of the study, developed the methodology, performed the majority of the investigations, conducted formal analysis, created the visualizations, and wrote the original draft. S.R. assisted with investigation and validation. Z.Z. contributed to the investigation. K.P. contributed to formal analysis and review and editing of the manuscript. V.S. assisted with methodology. K.C. contributed to conceptualization, managed the project, was responsible for project administration and funding acquisition, and contributed to writing, review, and editing.

## Supplemental Figure Legends

**Figure S1.**
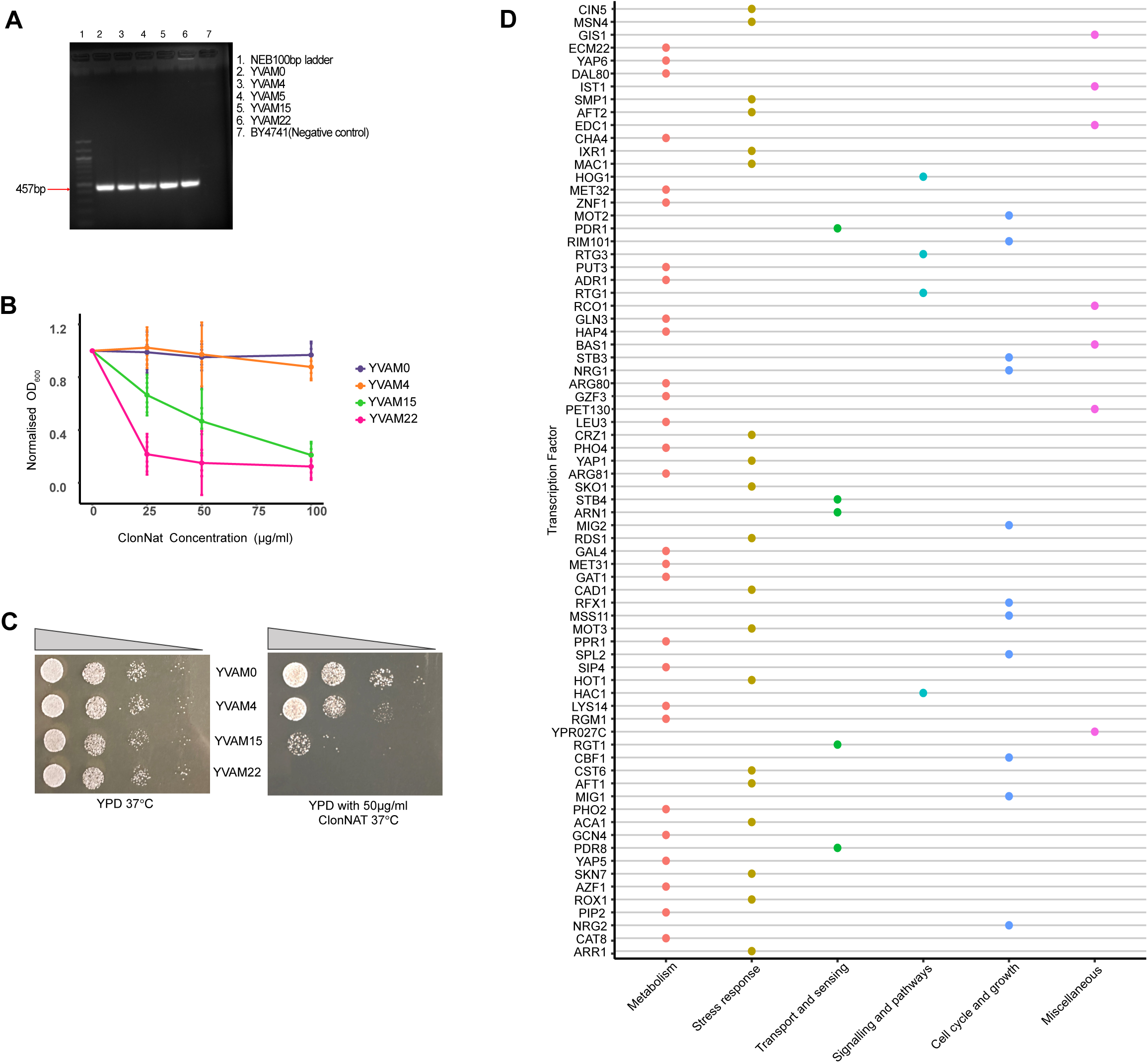
Characterization of NAT-integrated strains at elevated temperature. A. PCR confirmation of genome integration of NAT mutants. B. Growth of NAT integrated strains was assessed in YPD liquid medium supplemented with increasing concentrations of ClonNAT (0, 25, 50, and 100 µg/ml) at 37°C by taking optical density (OD) measurements at 600_nm_ after 12-16hrs (n=3). C. Spot dilution assay was performed on YPD agar plates with the indicated ClonNAT concentrations (50µg/ml) using 10-fold serial dilutions of each strain grown at 37°C for 48 hours (n=3). D. Bubble plot for functional categories of all the screened TFs. Y axis denotes transcription factors; X axis represents functional categories.

**Figure S2:**
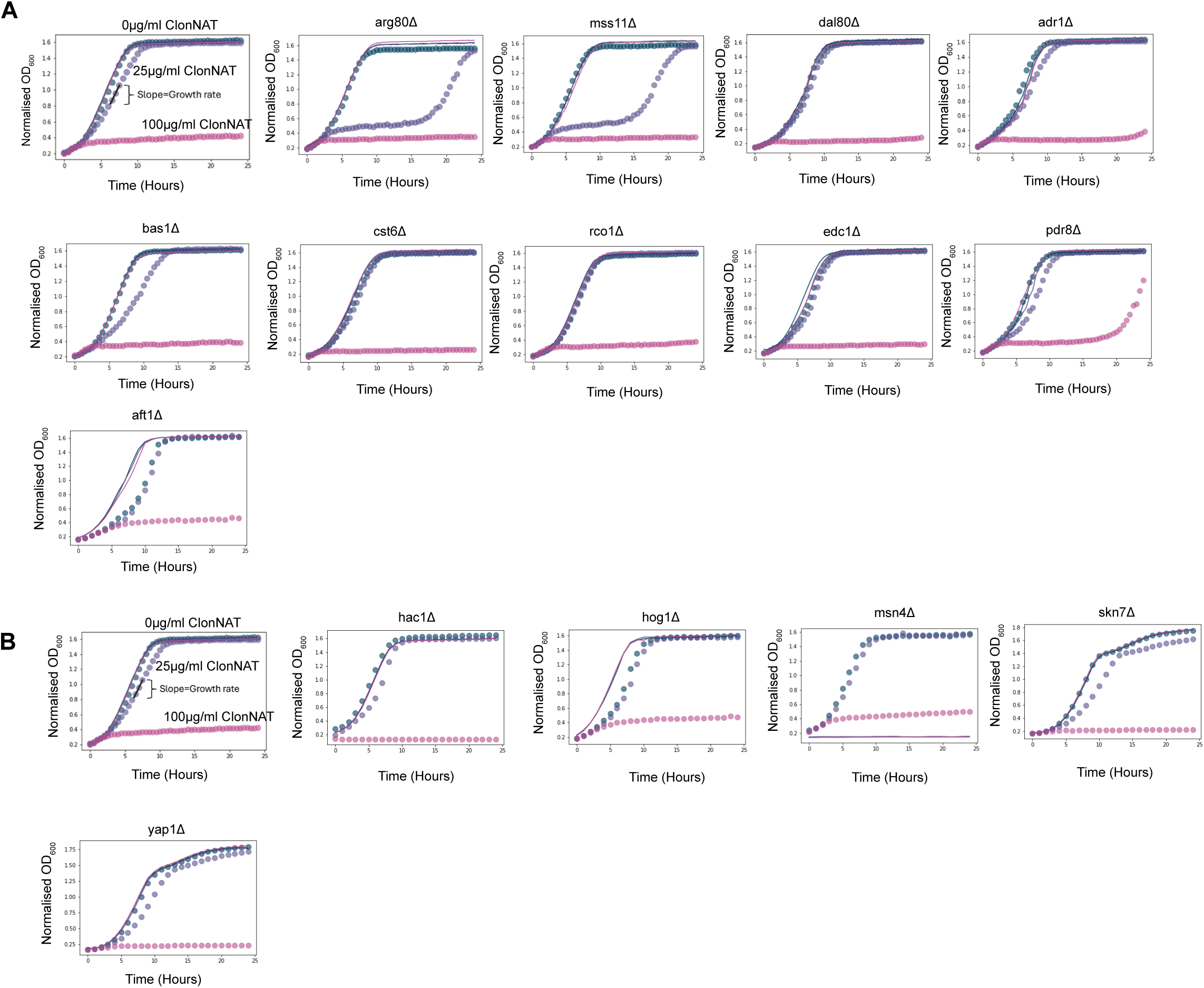
Growth kinetics of selected transcription factor deletion strains. A. Bioscreen growth curves of all TF hits from quadrants which had deletions having slower growth rate and longer time to reach the maximum growth (less M_max_ and more tM_max_). B. Bioscreen growth curves of all canonical TF screened.

**Figure S3:**
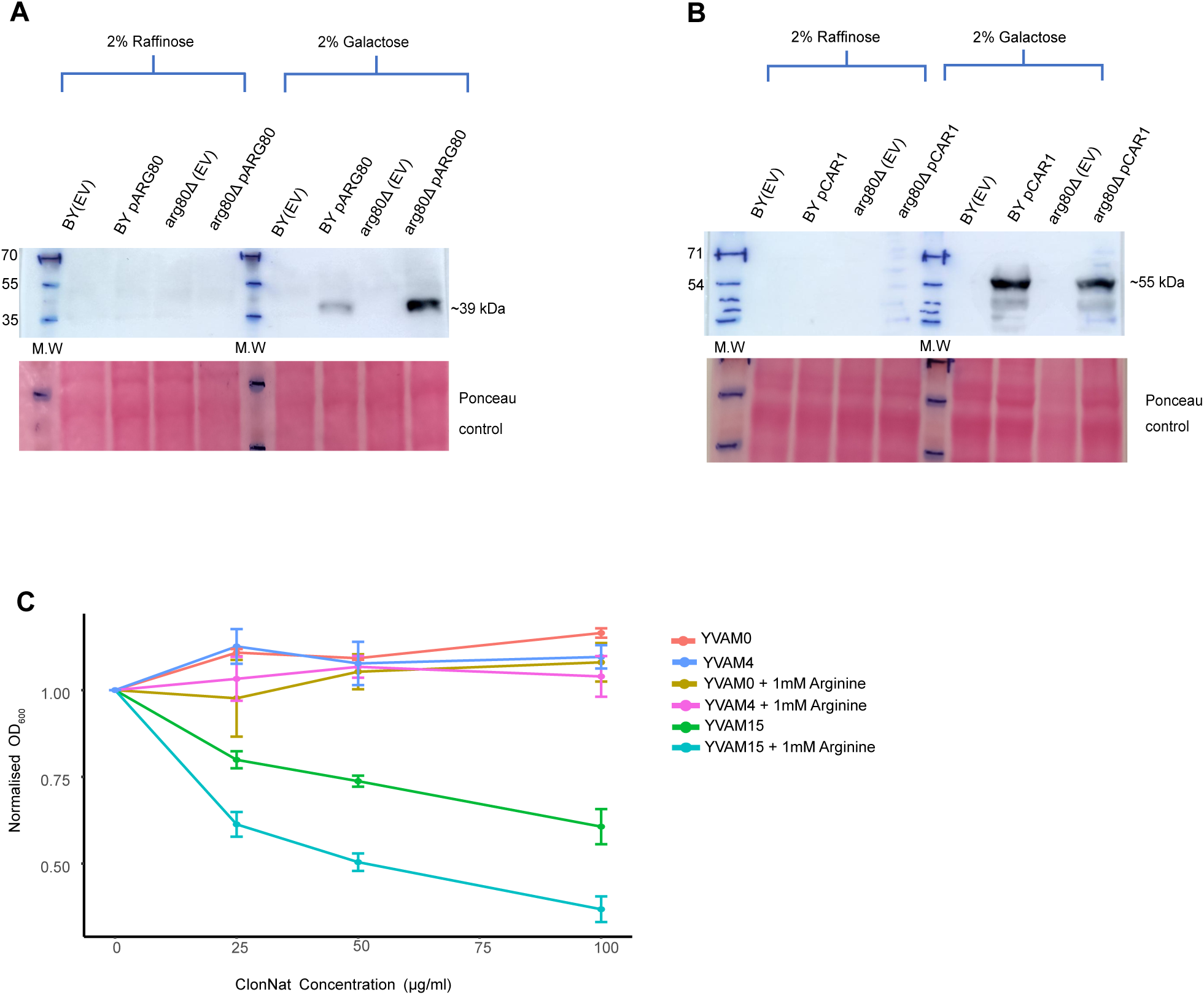
Validation of protein overexpression and effect of exogenous arginine on proteostasis. A. Western blot probed for His tag in ARG80 protein overexpressed (pARG80) via plasmid in BY4741 and arg80Δ with empty vector as control (EV) grown in 2% galactose (molecular weight ∼39 kDa) Shown here with ponceau as loading control (n=3). B. Western blot probed for His tag in CAR1 protein overexpressed (pCAR1) via plasmid in BY4741 and arg80Δ with empty vector as control (EV) grown in 2% galactose (molecular weight ∼55 kDa) Shown here with ponceau as loading control (n=3). C. Growth of YVAM0, YVAM4 and YVAM15 was assessed in SD media supplemented with increasing concentrations of ClonNAT (0, 25, 50, and 100 µg/ml) at 30°C with and without 1mM Arginine. by taking optical density (OD) measurements at 600_nm_ after 16hrs (n=3).

**Figure S4:**
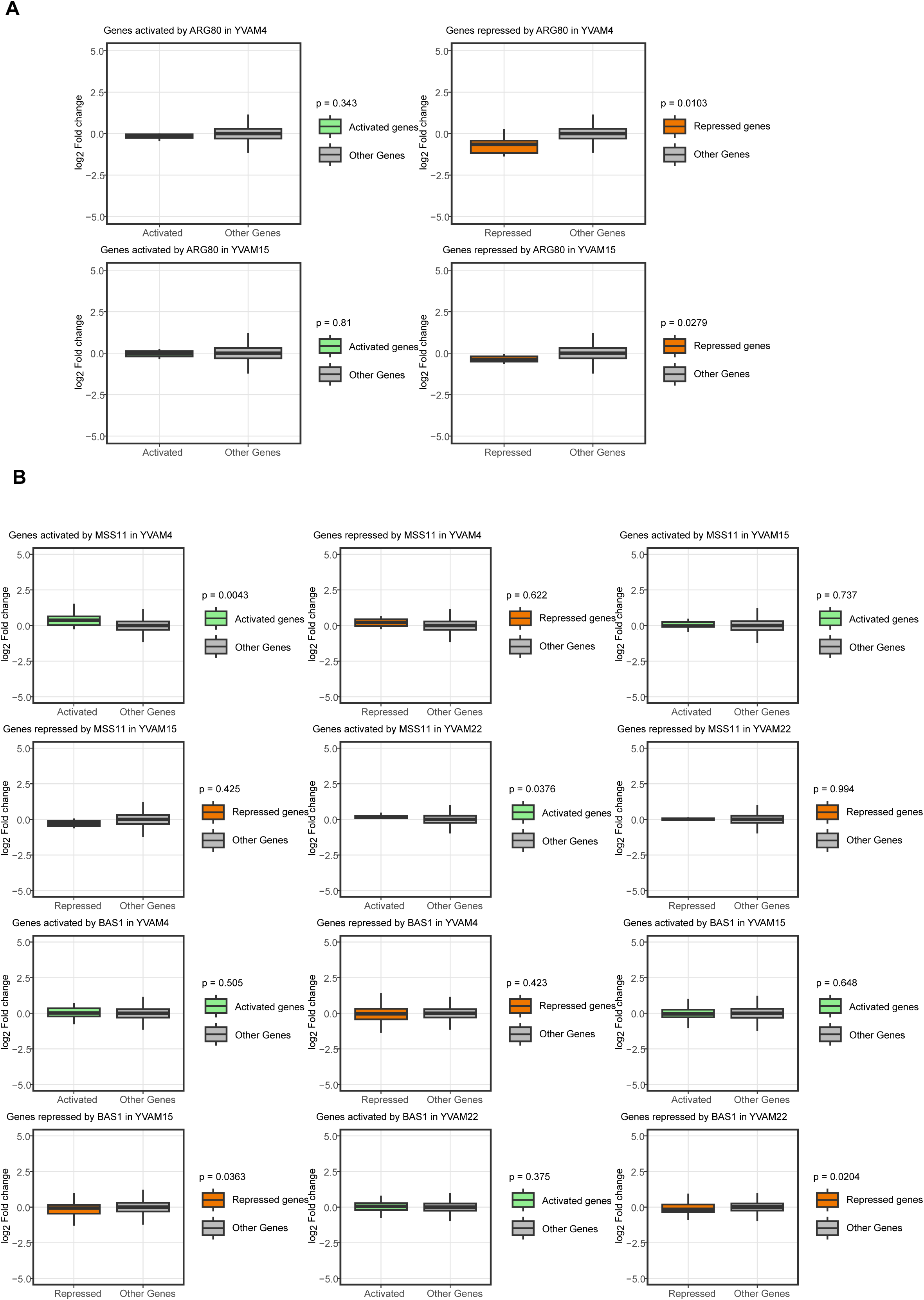
Transcriptional activity of Arg80, Mss11, and Bas1 in response to protein misfolding. A. Box plots show the distribution of log_2_ fold change (YVAM15 vs. YVAM0) and (YVAM4 vs. YVAM0) for genes activated by Arg80, genes repressed by Arg80, and all other genes (Rest). B. Box plots show the distribution of log_2_ fold change in presence of TS mutants for genes activated by Mss11 or Bas1, genes repressed by Mss11 or Bas1, and all other genes (Rest).

